# Metagenomics for bacterial spot pathogen and virulence factor tracking for Ohio fresh market tomato and pepper production

**DOI:** 10.64898/2026.05.01.717695

**Authors:** H. Toth, T.L. Klass, V. Roman-Reyna, F. Rotondo, D.M. Francis, M.T. Rodriguez, S.A. Miller, J.M. Jacobs

## Abstract

Bacterial spot is a consistent threat to global tomato and pepper productions; however, Ohio’s fresh market production currently lacks the updated surveillance data necessary to provide accurate management solutions. While traditional diagnostics focus on identification of a single causal agent, shotgun metagenomic sequencing (MGS) offers a comprehensive view of the infection court. An assignment-first MGS workflow was developed and validated in this study, utilizing Kraken2 databases to extract *Xanthomonas* species associated with bacterial spot and to characterize the microbial communities of bacterial spot in Ohio production systems. Through in silico spiking experiments, thresholds were established for bacterial spot identification. Species and pathovar identification via average nucleotide identity (ANI) remained accurate at abundance as low as 0.1%. A minimum of 2% *Xanthomonas* reads were required for high genome completeness (BUSCO >90%) and 3% for reliable type III secretion system (T3SS) effector profiling. Analysis of 63 samples from fresh-market production fields identified *Xanthomonas hortorum* pv. *gardneri*, *Xanthomonas euvesicatoria* pv. *euvesicatoria*, and *Xanthomonas arboricola* residing in symptomatic samples, alongside other taxa including Pseudomonas and Stenotrophomonas. Phylogenetic comparisons of metagenome-assembled genomes (MAGs) were comparable to whole genome sequences (WGS) from the same samples, supporting the reliability of culture-independent diagnostics. These results provide a robust framework for utilizing metagenomics as a diagnostic tool, expanding our knowledge of bacterial spot population structure in Ohio, and uncovering the bacterial communities associated with bacterial spot.

## Introduction

Tomato and pepper are significant crops grown throughout the world, with an estimated production of 186 million tons over 4.9 million hectares of tomato and 37 million tons over 2 million hectares of pepper in 2022 (2022). Production is threatened by bacterial spot, which is endemic to many areas where tomato and pepper are grown (Potnis et al. 2015). Bacterial spot of tomato and pepper are caused by as many as four *Xanthomonas* species: *X. euvesicatoria* pv. *perforans* (*Xep*), *X. euvesicatoria* pv. *euvesicatoria* (*Xee*), *X. hortorum* pv. *gardneri* (*Xhg*), and *X. vesicatoria* (*Xv*) (Bhattarai et al. 2017; Constantin et al. 2016; Jones et al. 2004). These xanthomonads have high genome plasticity, supporting their ability to adapt and evolve quickly (Jibrin et al. 2018; Thieme et al. 2005; Timilsina et al. 2015, 2019). Furthermore, bacterial spot species rely on the type III secretion system (T3SS) to secrete effectors into their host, contributing to pathogen virulence (Grant et al. 2006). Since species/pathovars cannot be differentiated by symptoms, current bacterial spot diagnostics rely on molecular techniques, such as conventional polymerase chain reaction (PCR) and quantitative PCR (qPCR). Morretti et al. (2009) created PCR primers to detect *Xanthomonas euvesicatoria* and was followed by the development of conventional multiplex PCR primers created by Araújo et al. (2012) to differentiate all *Xanthomonas* bacterial spot species and pathovars, but both methods can be time consuming, relying on gel electrophoresis. To resolve these labor-intensive strategies, Strayer et al. (2016) developed a qPCR assay targeting the *hrpB* region, enabling gel-independent differentiation of bacterial spot *Xanthomonas*.

Diagnostics are a front-line tool in the management of plant disease. Historically, plant disease diagnostics have relied on conventional, culture-based or molecular techniques, but these valuable approaches are limited as they identify a single agent. Bacterial spot has typically been described based on individual *Xanthomonas* species causing the leaf or fruit lesions that are characteristic of the disease. However, the biological reality is that these lesions contain a community of microbes that are part of the plant microbiome. The plant microbiome is a complex environment harboring beneficial, commensal and pathogenic microorganisms (Trivedi et al. 2020). During disease development, plant pathogens become dominant community members disrupting health (Malacrinò et al. 2022). Since many plant diseases have been exclusively associated with a singular causal agent, association or synergy with other microbes is understudied (Lamichhane and Venturi 2015).

A microbiome based diagnostic approach offers the opportunity to define all the organisms present in the infection court that may or may not participate in disease development. Despite the ongoing need for long-established culture-based techniques, next generation sequencing technologies provide an abundance of nucleotide-based information to both researchers and growers beyond traditional methods of diagnostics (Nizamani et al. 2023). Shotgun metagenomic sequencing (MGS) provides a unique lens of the total genetic content by providing both taxonomic assignment and functional potential of sequenced samples. While the microbial community composition can be a product of environmental, host, or microbe-microbe interactions (Trivedi et al. 2021) leveraging functional insights makes MGS a comprehensive plant pathogen surveillance tool. For example, MGS has proven to be powerful in instances such as low pathogen titer, culture-independent identification, and strain variation (Newberry et al. 2020; Román-Reyna et al. 2021; Yang et al. 2022).

MGS workflows for plant disease research can vary due to the experimental question or design being tested (Román-Reyna and Crandall 2024). For example, in the case of using metagenomics to test *Calonectria pseudonaviculata* detection on boxwood (Yang et al. 2022), blastn and Kraken2 read assignment were compared with custom fungal databases due to poor representation of fungal genomes in reference databases. In a study from our team, low abundance of *Xyllela fastidosa* was overcome by assembling reads assigned by Kraken2 (Román-Reyna et al. 2021).

Here, we used bacterial spot of fresh market tomato and pepper in Ohio as a model to test an assignment first approach for *Xanthomonas*. We employed an assignment first approach to analyze bacterial spot field samples from a state-wide survey in Ohio. We validated this method with an *in silico* experiment in addition to using it to test our field samples. Our work defined thresholds for *Xanthomonas* read requirement for metagenome-assembled genomes (MAGs) completeness, identification via ANI, and virulence factor presence. Furthermore, we investigated the bacterial communities associated with bacterial spot of fresh market tomato and pepper. Lastly, we compared MAGs developed from our assignment first approach with whole genomes from the same samples. Overall, we anticipate this work will lay the foundation for using metagenomic sequencing as a diagnostic tool, uncover microbial communities associated with bacterial spot of fresh market tomato and pepper, and expand knowledge of the bacterial spot type III secretion system effector repertoire.

## Materials and Methods

### Sample collection processing

A survey focused on bacterial spot of fresh market tomato and pepper of Ohio was conducted during the summer and fall of 2022. Three symptomatic leaves were collected from each variety present at sampling locations, and if present, one symptomatic fruit was collected per variety as well. We also randomly collected several asymptomatic tomato and pepper leaves from two of the farms. A 1-hole paper puncher was used to take 6-mm diameter tissue disks, centered on bacterial lesions, from leaves and fruit. The instrument was disinfested between each sample by soaking in 100% ethanol for 30 seconds. One disk was used for bacterial isolation, while two disks were used for DNA extraction and metagenomic sequencing.

### Bacterial isolation and identification

For bacterial isolation,400 µL of ultra-pure sterile water and a sterile metal bead were added to a 2.0 mL microtube and macerated using the QIAGEN TissueLyser II (QIAGEN, Venlo, NL: Cat. No. / ID: 85300) and the QIAGEN TissueLyser Adapter Set 2 x 24 (QIAGEN, Cat. No. / ID: 69982) at a speed of 30 Hz for two, one-minute intervals. After maceration, 20 µL of the macerate was transferred to a modified Tween Medium B (mTMB) plate using filtered pipette tips. The macerate was spread across the plate using a three-way streak using sterile loops. Plates were incubated at 28° C for 2-5 days, and single colonies with *Xanthomonas*-like features (slight to pronounced yellow pigmentation with a circular shape and convex elevation) were selected and streaked onto nutrient agar (NA) for additional purification. Single-colony isolates were stored in 1 mL of 25% glycerol for long term storage in our -80° C freezer. Immediately after streaking the macerate, 200 µL of Agdia ImmunoStrip^®^ (Agdia, Elkhart, IN: ISK 14600/0025) buffer was added to the remaining macerate and a *Xanthomonas* genus ImmunoStrip^®^ was dipped in the solution. We proceeded only with positive samples.

For further *Xanthomonas* identification, we conducted a colony PCR using primers based on the *gyrB* gene (Parkinson et al. 2007). Template DNA for each sample was created by placing a 1 µL loopful of bacteria into 40 µL of a TE-T solution, mixing the solution using a vortex at speed 10 for 15 seconds, incubating the tubes at 95° C for 5 minutes, and spinning them down for 5 minutes. Using the recommendation for the Quick-Load *Taq* 2x Master Mix (New England Biolabs, Ipswich, MA: Cat. No. M0271L) 25 µL reaction, we used 2 µL template DNA, 1 µL forward primer, 1 µL reverse primer, 12.5 µL Quick-Load *Taq* 2x Master Mix, and 8.5 µL ultra-pure sterile water. For positive controls, template DNA from known *Xanthomonas* strains was used. Negative control was ultrapure water. The following program was run on the thermocycler for all samples: 2 minutes initial denaturation at 95° C, 35 cycles of 30 seconds at 95° C, 30 seconds at 50° C, and 30 seconds at 68° C, and a final extension for 5 minutes at 68° C. Results were visualized by loading 10 µL of each product into a 1% agarose gel stained with SYBR Green I Nucleic Acid Gel Strain at 10,000x (Fisher Scientific, Waltham, MA: Cat. No. S33102) at 100V for 60 minutes. A blue LED transilluminator was used to visualize the gel, and samples that produced a band of 750 base pairs were considered to belong to the genus *Xanthomonas*.

### DNA extraction

To prepare for DNA extraction, *Xanthomonas* isolates were grown on NA plates at 28° C for 2-3 days. A 10 µL inoculating loopful of bacteria was added to 100 µL of ultra-pure sterile water and DNA was extracted using the Monarch Genomic DNA Purification Kit (New England Biolabs, Cat. No. T3010S), following the protocol for cultured cells. After DNA extraction and elution, the concentration was measured with a spectrophotometer.

Total DNA for metagenomic sequencing was extracted using, using the QIAGEN DNeasy Plant Pro Kit (QIAGEN, Cat. No. / ID: 69204) following the manufacturer’s protocol with slight modifications. Sterile, 1000 µL pipette tips were used to transfer tissue disks into the QIAGEN Tissue Disruption Tubes. Modifications include macerating the samples at a speed of 30 Hz, adding 30 µL of Buffer EB, and incubating the tubes at 37° C for 10 minutes before elution. After DNA extraction and elution, the concentration was measured with a spectrophotometer.

### Whole genome and metagenomic sequencing

All sequencing was conducted at the Infectious Disease Institute Genomics and Microbiology Solutions Laboratory (IDI-GEMS) at The Ohio State University. For both whole genome and metagenomic sequencing, 2 µL of DNA was used as input for sequencing library preparation. The Illumina Nextera XT DNA Library Prep kit, which specifically used the Integrated DNA Technologies (IDT) for Illumina DNA UD Indexes. At IDI-GEMS, both whole genome and metagenomic samples were sequenced on Illumina’s NextSeq 2000 at a read length of 150 base pairs (bp) x 2. Whole genomes were sequenced with a goal of obtaining 1-2 million reads per sample. The majority of metagenomic samples were sequenced with a goal of 12.5 million reads per sample, while a select few were sequenced with a goal of 25 million reads per sample.

### MAG and whole genome assembly

We created a metagenomic workflow for identification and analysis of samples suspected to be infected with *Xanthomonas*. Our pipeline centers around using Kraken2 (Wood et al., 2019) taxonomic assignment and extraction of taxon specific reads to create MAGs, which we term an “assignment-first” approach. The database we define as “standard” is the Kraken2 plus-pf database (Figure 4B). Our database was made specifically for bacterial spot identification. It contains an equal number of genomes (n=33) for each species and pathovar in the bacterial spot disease complex as well as *X. arboricola* genomes. Furthermore, *Stenotrophomonas* (n=33) and plant genomes (n=2) were also included in this database to avoid mis-assignment of reads.

Comparison of standard and databases revealed very few differences in *Xanthomonas* reads, therefore, we chose to use MAGs created from the database for all analyses.

Prior to taxonomic classification, all metagenomic data was trimmed using Trimmomatic, where the first 15 bases and any bases beyond 150 bases were trimmed from each read (Bolger et al. 2014). After taxonomic classification with Kraken2, KrakenTools, extract_kraken tool (Lu et al. 2022), was used to extract *Xanthomonas* reads (NCBI taxid = 338) from Kraken2 output.

Extracted reads were assembled into MAGs using metaSPAdes (v. 3.15.4) (Nurk et al. 2017). BUSCO was used to determine the percent completeness of assembled MAGs, specifically using the xanthomondales_10db database (Manni et al. 2021).These steps were repeated for every metagenomic sample, including those in our *in silico* analyses.

For whole genome data, the first 15 bases and any bases beyond 150 were trimmed from each read using Trimmomatic (v. 0.40) (Bolger et al., 2014). Unpaired reads were concatenated to be used in assembly. Trimmed and unpaired reads were assembled into a single genome using Unicycler (v. 0.5.0) (Wick et al. 2017). Contigs less than 200 bp were deleted. Completeness of whole genomes was determined by BUSCO (v. 5.0.0), using the xanthomonodales_10db database (Manni et al. 2021).

### Metagenomic analysis

We conducted an *in silico* experiment to test our workflow, where we created metagenomic samples by spiking *Xee* whole genome reads (OSU1673) into asymptomatic metagenomic pepper reads. Samples contained between 0.05%-4% *Xanthomonas* reads, which were chosen based on observations made in this study. We used asymptomatic metagenomic pepper reads to mimic the phyllospheric microbiome of an Ohio fresh market production system. Seqtk was used to subset reads randomly from both the asymptomatic metagenomic pepper and the whole genome reads, which resulted in a total of 12.5 million reads per sample, regardless of ratio (Li 2013). The subset process was repeated five times with five different seeds chosen with a random number generator. After subsetting, pepper and whole genome reads were concatenated, and the resulting paired reads were put through our metagenomic workflow.

All ANI and genome coverage calculations were calculated by pyani, using default parameters (Pritchard et al. 2015). R (v. 4.4.0) (R Core Team 2024) and the ComplexHeatmap package (Gu 2022; Gu et al. 2016) was used to create heatmaps and dendrograms from pyani outputs. ANI calculation method was ANIm, and genome coverage was calculated by the following formula: (read length x # of reads) / haploid genome length. Reference genome accession can be found in Supplemental Table 1.

Taxonomy abundance graphs were created with data derived from Kraken2 assignment from the plus-pf database. Kraken-biom was used to transform Kraken2’s kreport outputs into a BIOM format (Dabdoub 2016). The graphs were created in R, using the following libraries: readxl (Wickham and Bryan 2025), phyloseq (McMurdie and Holmes 2013), microbiome (Lahti and Shetty 2017), vegan (Oksanen et al. 2025), and tidyverse (Wickham et al. 2019). In conjunction with these packages, custom R scripts developed by the Research and Analytical Service Cores (RASC) at the Ohio State University (Wooster, OH) were used to visualize taxonomy abundance graphs. We filtered all bacterium reads that made up less than 1% of the total to be classified under “other (rare)”.

Both Non-Metric Multidimensional Scaling (NMDS) and Shannon diversity calculations were performed in R. NMDS was preformed using Bray-Curtis distances using phyloseq’s ordinate() function. Significance was assessed with PERMANOVA via vegan’s adonis2() function.

Shannon diversity was calculated using vegan’s diversity() function, using the index “Shannon.” The Kruskal-Wallis test was used to calculate significance among host and region Shannon diversity values.

All MAGs, corresponding whole genomes, and whole genome references (Figure 5A) were run with roary (Page et al. 2015). Roary’s core genome alignment output was used as input for mycotools’ fa2tree tool, which uses IQtree to create phylogenetic trees (Konkel and Slot 2023). Bootstrap value of 1000 was used in tree creation.

To determine the presence and absence of type III secretion system (T3SS) effectors, we created a custom protein blast database. We used the EuroXanth web resource as a reference for our database creation (euroxanth.ipn.pt/doku.php?id=bacteria:t3e:t3e). This resource was consulted while curating *Xanthomonas* specific protein sequences from UniProt (The UniProt Consortium 2023). All protein sequences were concatenated into a singular fasta file to make into a BLAST database, where makeblastdb was used to create our database (Camacho et al. 2009). Local blastx was used to compare all genomes to our custom database (Camacho et al. 2009). Presence of an effector was determined if the following two thresholds were met: ≥ 70% percent identity and ≥ 50% coverage, where coverage was calculated by length/slen.

### Grower perception survey

Diagnostic tools are only as good as the end-user deems them. Therefore, to evaluate grower perceptions of using sequencing technologies for diagnostics and surveillance of bacterial spot, we developed a survey for growers to take when samples were collected. The term “Ohio PathID” was used to define using sequencing technology for diagnostics and surveillance to make the concept more accessible. We used the diffusion of innovations (DOI) model as our framework based on models used for related research (Rogers 2003). The DOI constructs relevant for this study include relative advantage, compatibility, complexity, and observability (Rogers 2003). Each construct had several questions in the survey (five for relative advantage and three for the others) that the growers were asked to respond to using a 5-point Likert-type scale from 1 (strongly disagree) to 5 (strongly agree). Although we visited 32 farms across Ohio, we only received usable survey responses from 24 farms. Our survey received IRB exemption on 7/26/2022 (IRB Protocol Number: 2022E0688).

## Results

### Development, implementation and evaluation of Ohio PathID Approach

To develop and field test a metagenomics-based approach for bacterial spot pathogen identification, we conducted a statewide survey of fresh market tomato and pepper production for this disease. Symptomatic tissue was collected from 32 open field production farms across 22 counties in Ohio resulting in a total of 63 samples, split between 29 tomato, 26 pepper and 8 weedy hosts (Figure 1A). At the same time of symptomatic tissue collection, qualitative data was gathered on farmer perceptions of our diagnostic approaches. To ensure these sequencing-based methodologies were of interest to the end stakeholder, we collected preliminary perception data from 24 growers on the use of sequencing technologies in plant pathogen diagnostics and surveillance (termed Ohio PathID). Grower perceptions of Ohio PathID were generally positive upon initial introduction (Figure 1B). Diffusion of innovation constructs, as defined by (Rogers 2003), were used to categorize survey responses. The constructs relative advantage, compatibility and complexity survey responses ranged from neutral to strongly agree.

**Figure 1.**
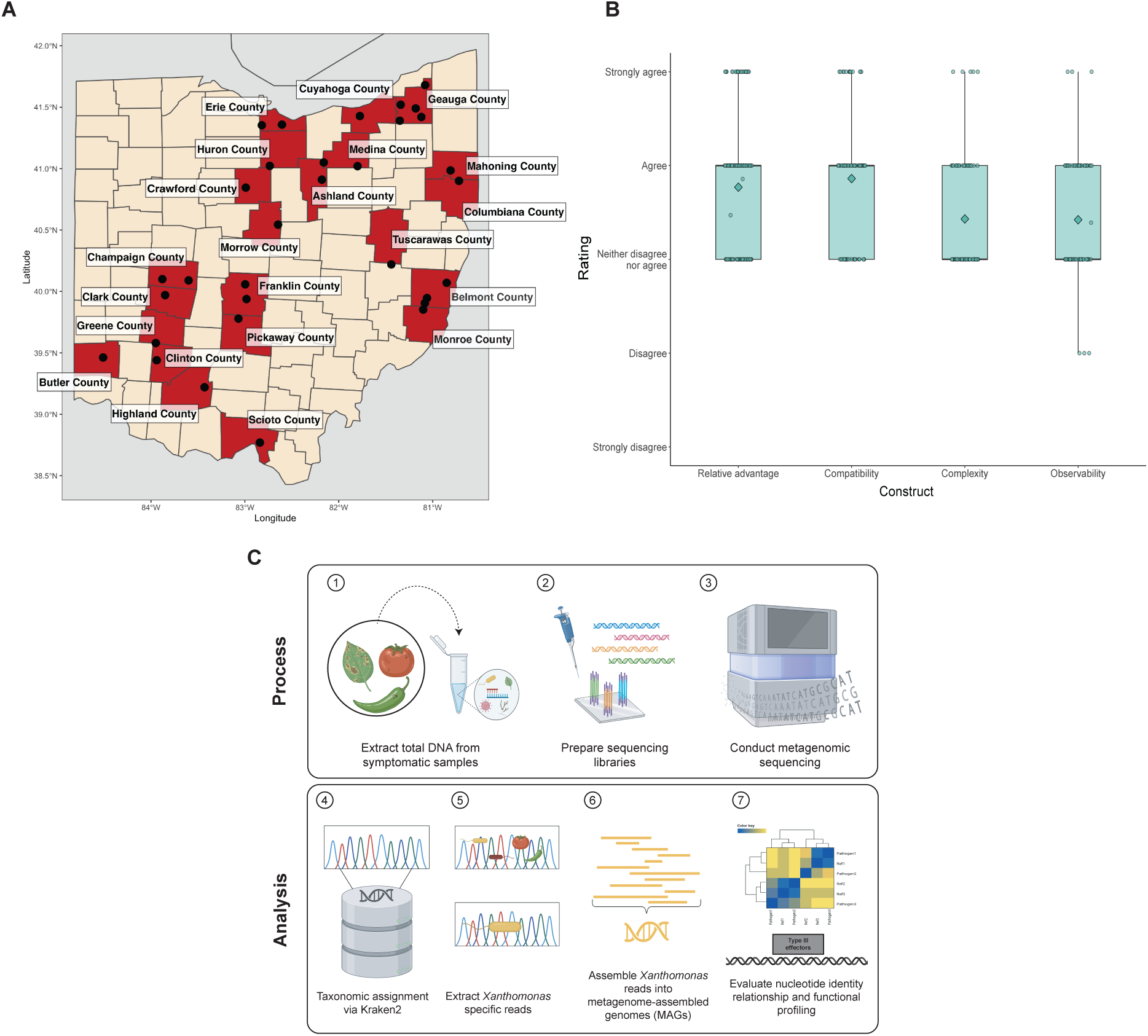
Map of sample collection, grower perception survey results and workflow for processing and analysis. **A)** Map of Ohio, where red highlights counties from which where samples were collected. Black dots show approximate location of each farm. **B)** Grower perceptions of genomic technology prior to tissue collection (n=24). Diffusion of innovation constructions are along the x-axis, and grower answers are represented by boxplots. Individual data points represent a single response, and diamonds represent the mean for that construct. **C)** Workflow for processing, sequencing, and analyzing metagenomic data from Ohio fresh market tomato and pepper samples.

Observability was the only construct with disagreement responses (n = 3).

### In silico experiment provides foundational thresholds for Xanthomonas detection via MGS

We used an assignment-first workflow approach for the detection of bacterial spot in metagenomic data of fresh market tomato and pepper in Ohio (Figure 1C). To evaluate the relationship between *Xanthomonas* read abundance and MAGs, we conducted an *in silico* experiment where *Xee* whole genome reads were spiked into asymptomatic metagenomic pepper reads. Assessment of the experiment was determined by genome metrics such as average nucleotide identity (ANI), genome converge, type III secretion system (T3SS) effector identification and Benchmarking Universal Single-Copy Ortholog (BUSCO) score. We obtained a strong linear correlation between amount of *Xanthomonas* spiked in and the amount of *Xanthomonas* reads recovered from Kraken2 taxonomic assignment (percentage of *Xanthomonas* spiked in sample, y = 318 + 0.999x, R^2^ = 1.00) (Figure 2A). BUSCO scores for MAGs assembled from *Xee in silico*-spiked samples containing 4%, 3%, and 2.5% *Xanthomonas* reads scores were 98.6%, 95.5%, and 91.1%, respectively, which was close to the whole genome isolate (OSU 1673) BUSCO score of 99.3% (Figure 2C) (Supplemental table 2). In contrast,

**Figure 2.**
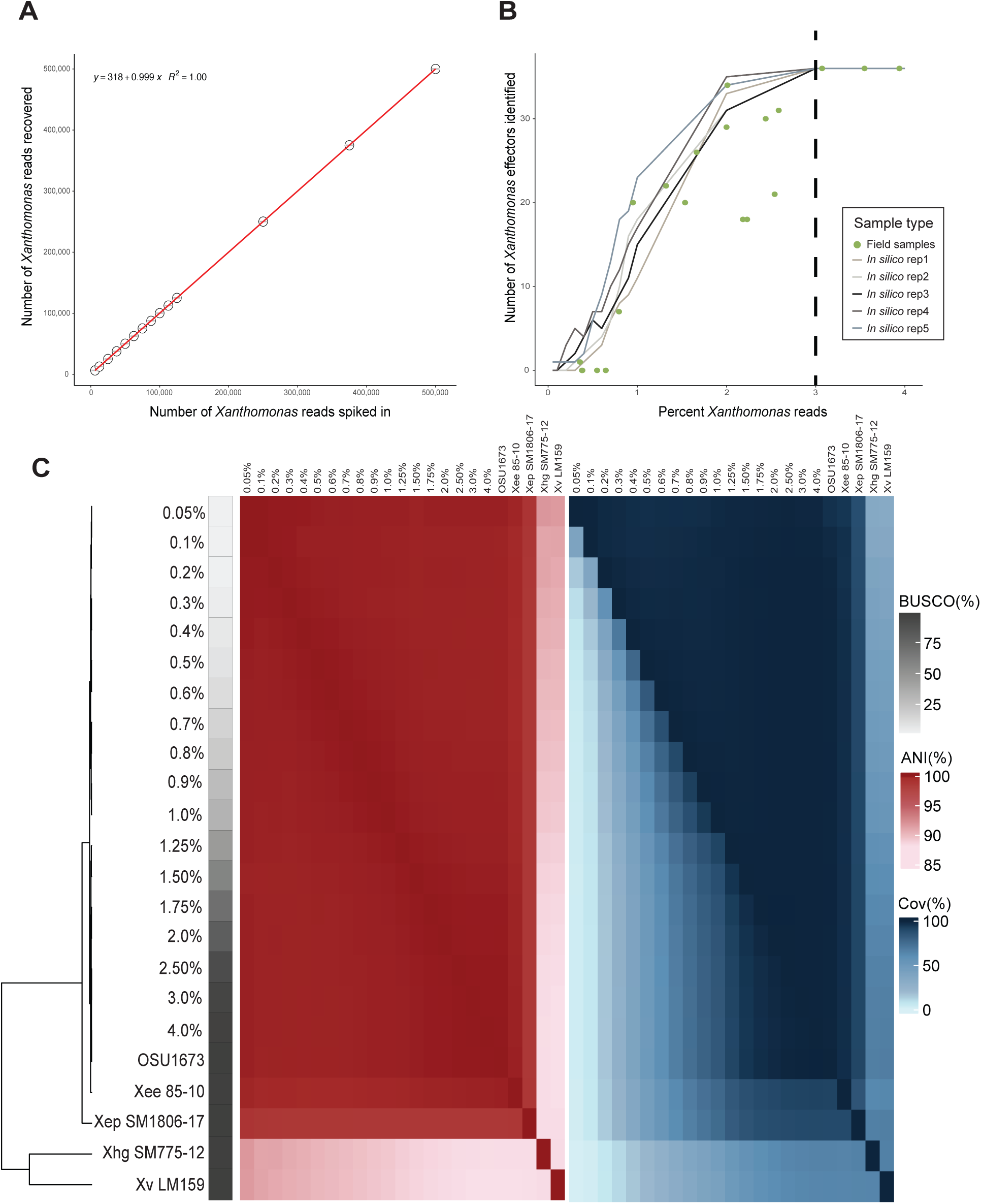
*In silico* validation of bioinformatic workflow. We spiked *in silico Xee* whole genome reads (OSU1673) into metagenomically sequenced asymptomatic pepper DNA to set thresholds needed for functional recovery. Spiking was conducted with seqtk. **(A)** *Xanthomonas* reads recovered by Kraken2 (y-axis) spiked in at incremental percentages (x-axis) with small red bars representing the standard error across replicates (n=5). **B)** *Xanthomonas* effectors identified by blastn between *in silico* samples (grey-scale, n=5) and field samples (green dots). Effectors were considered present if they had a percent identity above 70% and coverage above 50%. Coverage was calculating by dividing the length of the gene in the MAG (length) and the length of the gene in the database (slen). All field samples were sequenced at a depth of 12.5M reads. Dotted line indicates a hypothesized effector identification threshold. **C)** BUSCO score, average nucleotide identity (ANI) and coverage convey percentage of *Xanthomonas* reads for required for downstream analysis. Replicate one was used for panel C analyses. All reference genomes (*Xee* 85-10, *Xep* SM1806-17, *Xhg* SM775-12, and *Xv* LM159) were downloaded from NCBI (Supplemental Table 1).

MAGs from the samples containing 2% and 1% *Xanthomonas* reads had scores of 80.4% and 31.6%, respectively. All MAGs generated from samples with less than 1% *Xanthomonas* reads had BUSCO scores below 26.1% (Figure 2C). Hierarchical clustering based on ANI calculations exhibit MAGs with 1% *Xanthomonas* reads or less cluster separately than MAGs from samples with higher *Xanthomonas* read percentages. MAGs from samples with 1.25% of *Xanthomonas* reads or higher clustered together with the *Xee* reference (*Xee* 85-10). (Figure 2C).

Genome coverage between *Xee* 85-10 and MAGs varied widely, between 2.12%-91.77% (Figure 2C) (Supplemental Table 2). However, genome coverage between *Xee* 85-10 and MAGs does not drop below 90% until *Xanthomonas* reads are equal to or less than 1.75% (Supplemental Table 1). ANI stayed consistent for all MAGs, regardless of sample origin, as the MAGs coming from samples with lowest and highest percentage of *Xanthomonas* reads (0.05% and 4%) had ANI values of 99.73% and 99.87%, respectively. Despite BUSCO, clustering, and coverage suggesting a minimum threshold of 2% *Xanthomonas* reads for MAG identification, effector presence, with both *in silico* and field samples demonstrated otherwise. We found all *in silico* replicates and field samples identified 36 T3SS effectors when containing 3% and 4% *Xanthomonas* reads (Figure 2B). MAGs from *in silico* and field samples with less than 3% *Xanthomonas* reads recorded varying effector presence.

### Microbial diversity and Xanthomonas detection in Ohio Tomato and Pepper fields

We used 63 samples from our survey of fresh market tomato and pepper across Ohio (Figure 1B). After taxonomic assignment with Kraken2’s PlusPF database (Wood et al. 2019), we specifically looked at relative bacterial read abundance, and found a high relative abundance of *Xanthomonas*, *Pseudomonas*, *Stenotrophomonas*, and *Pasteurella* (Figure 3A). Out of 63 samples, we successfully isolated *Xanthomonas* from 23 of them (Figure 3A). However, the relative abundance nor the absolute amount of *Xanthomonas* reads in a sample did not necessarily correspond with successful isolation. For example, sample 81819 had 369,126 *Xanthomonas* reads and a relative abundance of *Xanthomonas* of 68%, but no *Xanthomonas* isolates were recovered (Supplemental Table 3 and 4). Conversely, samples 22Brandy and 32T2Fruit had 10,916 and 77,556 *Xanthomonas* reads and had a bacterial relative abundance of *Xanthomonas* of 0.097% and 0.018%, respectively (Supplemental Table 3 and 4). Yet, isolates were successfully recovered from both samples.

**Figure 3.**
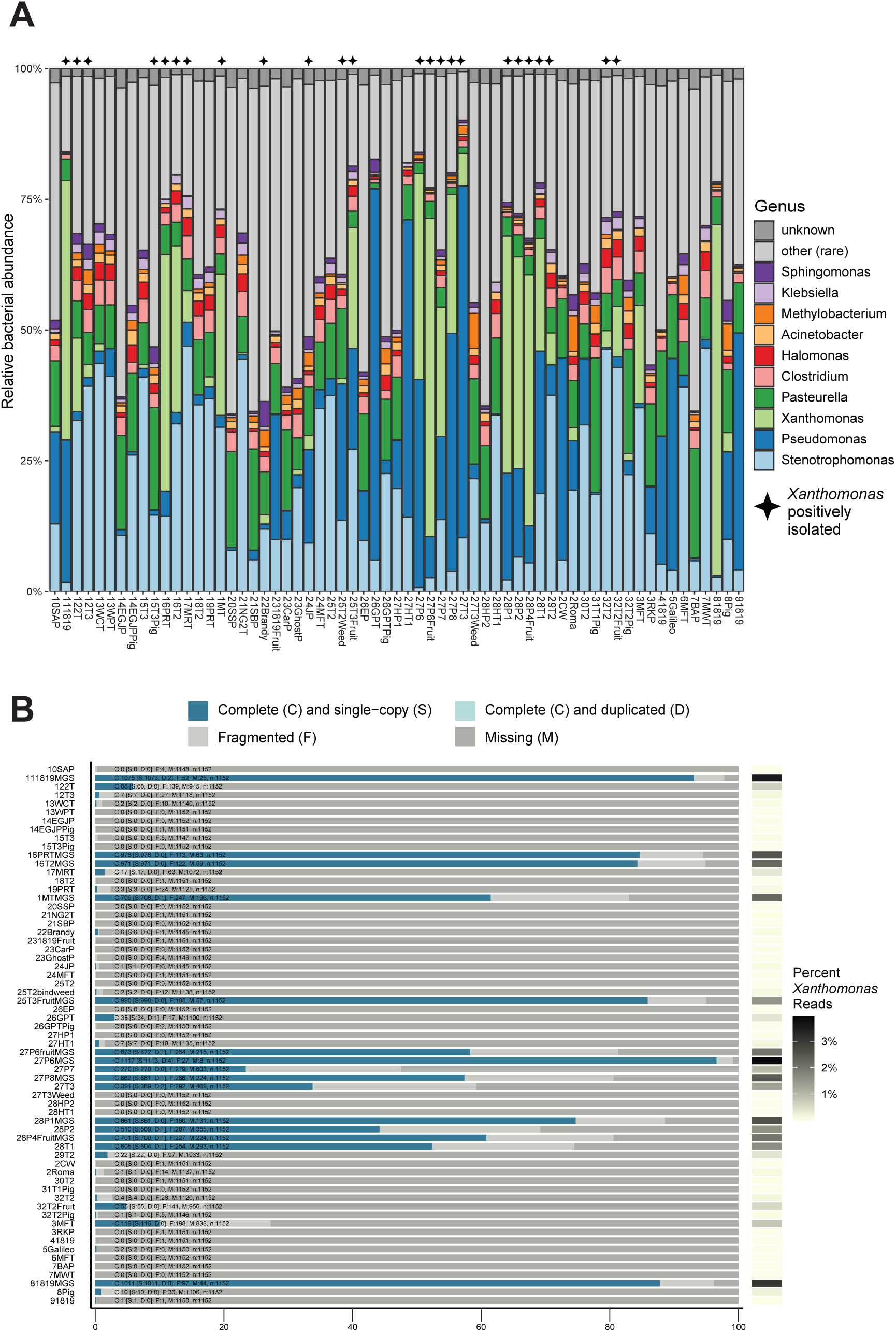
Bacterial spot pathogen identification thresholds. **A)** Kraken2 plus-pf database resolved bacterial taxonomic relative abundance in tomato and pepper samples at the genus-level. “Unknown” refers to reads that did not receive an assignment, and “other (rare)” corresponds to bacterium that constituted less than 1% of total bacterial reads. Star indicates that *Xanthomonas* was isolated and sequenced with Illumina. **B)** BUSCO score was compared to percent of *Xanthomonas* reads. BUSCO score was determined for each MAG. Dark blue and light blue bars indicate complete marker genes present, while the light and dark grey indicate fragmented or missing marker genes. Percent *Xanthomonas* reads is indicated by the heatmap on the right, with darker boxes indicating a higher percentage of *Xanthomonas* reads.

In samples where *Xanthomonas* bacterial relative abundance was greater than 5%, the species *X. hortorum, X. euvesicatoria* and *X*. *arboricola* were identified (Supplemental Table 3). In each of these samples, one *Xanthomonas* species dominated, although some samples did have multiple *Xanthomonas* species present. For example, in sample 1MT, *X. hortorum* had a relative abundance of 22.63%, whereas *X. arboricola* accounted for only 2.22%. In contrast, there was a larger variety of *Pseudomonas* species abundance across samples. In samples where *Pseudomonas* bacterial relative abundance was above 5%, we found *Pseudomonas viridiflava*, *syringae*, *fulva*, *rhizosphaerae*, *argentinensis*, and *psychrotolerans* (Supplemental Table 4).

Several samples contained multiple *Pseudomonas* species with similar relative read abundances. For example, sample 2CW had 6.37% and 6.28% relative abundance of *P. syringae* and *P. argentinensis*, respectively. Lastly, *Stenotrophomonas* and *Pasteurella* relative abundance were comprised of a singular species across all samples: *Stenotrophomonas maltophilia* and *Pasteurella mulocida*.

Alpha diversity, measured by the Shannon index, was not significantly different among hosts (tomato, pepper and weedy) (*P* = 0.79) or region (Central, east, northeast and northwest) (*P* = 0.72) (Supplemental Figure 1A and 1B). Beta diversity, measured by Bray-Curtis distances, was significantly different by host (R^2^ = 0.33, *P* > 0.001) but not region (R^2^ = 0.06, *P* > 0.214) (Supplemental Figure 1A and 1B).

We determined some field samples did not follow the same BUSCO score trend seen with *in silico* samples by comparing percentage of *Xanthomonas* reads in each sample with its corresponding BUSCO score. Samples 27P6, 111819, and 81819 were the only samples with greater than 3% *Xanthomonas* reads, (3.94%, 3.54%, and 3.07%, respectively) and their BUSCO scores reflect their high *Xanthomonas* content, with scores of 96.9%, 93.3%, and 87.7%, respectively (Figure 3B) (Supplemental Table 4). However, MAGs from samples with below 3% *Xanthomonas* reads do not necessarily follow this same trend. For example, sample 25T3Fruit had a BUSCO score of 85.9%, only having 1.54% *Xanthomonas* reads, while samples with between 2-3% *Xanthomonas* reads received BUSCO scores ranging from 57% to 84.5% (Figure 3B). Although our *in silico* experiment defined a 3% *Xanthomonas* read minimum for functional profiling, it showed tools such as ANI could be useful in identifying MAGs created from samples with a low abundance of *Xanthomonas*. ANI was calculated between MAGs developed from samples containing more than 0.10% *Xanthomonas* reads and reference xanthomonads.

Regardless of the percentage of *Xanthomonas* reads that a MAG came from, all MAGs had an ANI value greater than 95%, the cutoff for ANI species determination (Goris et al. 2007), with the reference genome it clustered with (Figure 4) (Supplemental Table 2).

**Figure 4.**
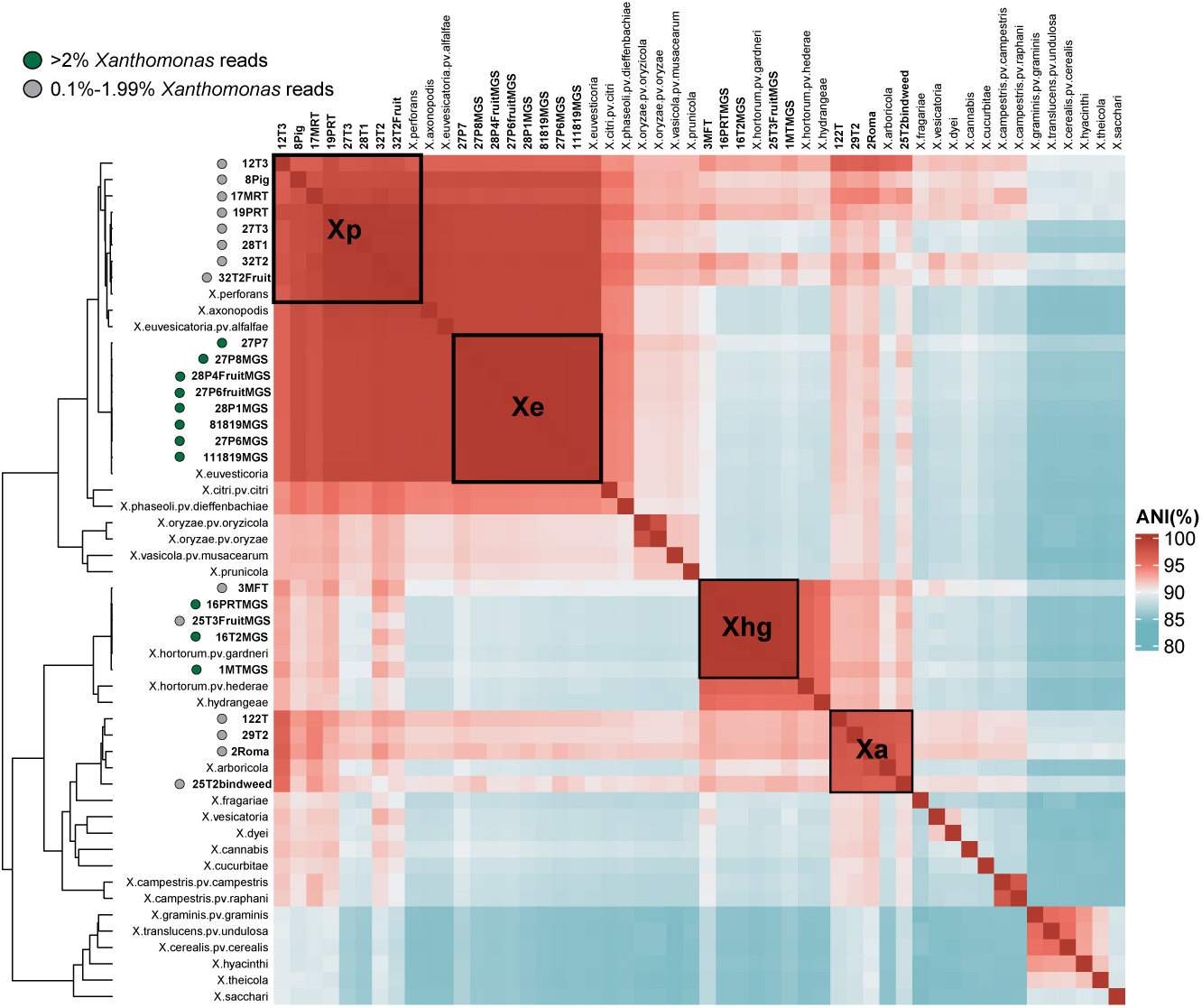
ANI-based identification of *Xanthomonas* from tomato and pepper metagenomes. ANI of all *Xanthomonas*-derived MAGs (bold), whole genome sequenced (black stars) and NCBI xanthomonad reference genomes. All MAGs (bold) are denoted with green or gray circles if Xanthomonas reads were greater that 2% or between 0.10% to 1.99%, respectively. NCBI references defined in Supplemental Table 4.

### Comparison of MAGs to their respective whole genome sequences

To determine if MAGs are comparable to whole genomes in a phylogenetic analysis, we created a core gene phylogeny of MAGs, whole genome sequenced (WGS) isolates from the same sample, and appropriate bacterial spot *Xanthomonas* reference genomes. We only used MAGs from field samples with BUSCO scores greater than 74.6% and contained greater than 2% *Xanthomonas* reads. Each MAG and WGS sample clustered next to each other in our phylogeny, where they were supported by a 100% (1000/1000) bootstrap value (Figure 5A). T3SS effector identification between MAGs and WGS pairs was consistent with a few exceptions (Figure 5B). The OSU65/28P1MGS pair was the only pairing where the WGS contained all effectors identified between the two and contained additional unique effectors. 28P1MGS harbored 31 effectors, all shared with OSU65, while OSU65 possessed five unique effectors, totaling at 36 effectors. In contrast, OSU64/27P6MGS and OSU24/111819MGS had a singular instance where the MAG identified one more effector than the corresponding whole genome. 27P6MGS contained effector *xopD* while it was absent in OSU64, and 111819MGS had effector *xopE2*, while it was absent in OSU24. The remaining pairs, OSU34/16PRTMGS and OSU35/16T2MGS, presented shared and unique effectors for both the MAG and WGS. OSU34 and 16PRTMGS had a shared twenty effectors, while OSU34 had effector *avrBs3* and 16PRTMGS had effector *xopH1*. OSU35 and 16T2MGS had a shared 18 effectors, while OSU35 had effectors *xopAM* and *xopK* and 16T2MGS had effector *xopH1*.

**Figure 5.**
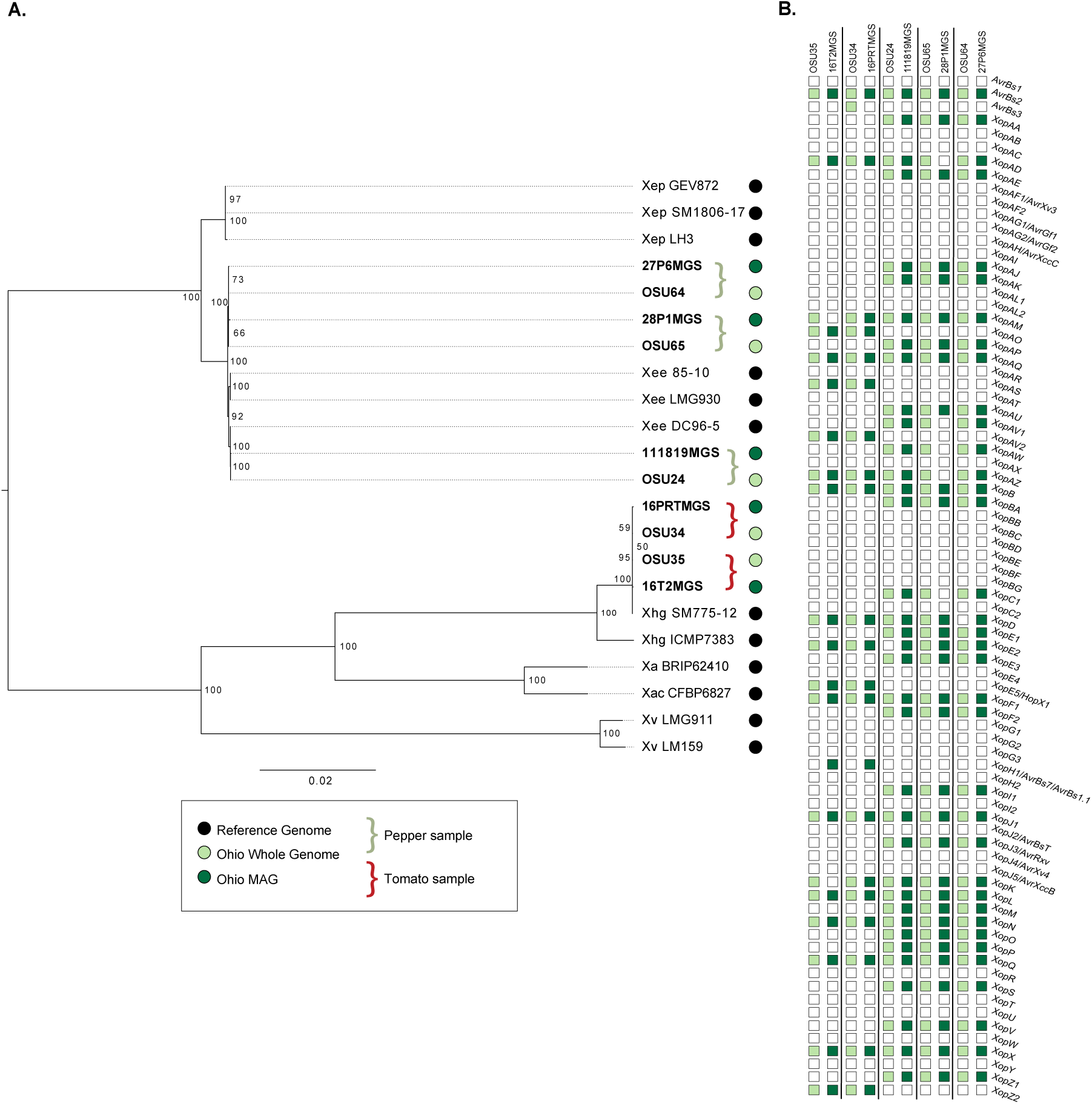
Whole genome sequenced isolates and MAGs from the same sample are nearly identical. **A)** Core genome tree of Ohio tomato and pepper-derived MAGs (dark green), Ohio whole genomes (light green), and publicly available reference genomes of bacterial leaf spot xanthomonads (black) (Supplemental Table 1). Curly brackets indicate the same Ohio sample, while bracket color indicates sample host (e.g. pepper, green; tomato, red) of isolation. Tree was created using the core genome alignment produced by roary and the fa2tree tool in mycotools, which uses IQtree (bootstrap value = 1000). **B)** T3SS effector repertoire of MAGs and their corresponding whole genome isolate. Green-filled squares represent the effector was identified (MAGs, dark green; whole genomes, light green).

## Discussion

In this study, we conducted a state-wide survey of bacterial spot from fresh market tomato and pepper production in Ohio to test disease detection through metagenomic sequencing. This was motivated by positive grower perceptions to surveillance technologies, such as genomics and metagenomics.

We were successful at identifying *Xanthomonas* bacterial spot pathogens with metagenomics. By pairing a bacterial spot-centered database with Kraken2, we could retrieve *Xanthomonas* reads from every sample, regardless of abundance. This allowed us to understand the relationship between amount of *Xanthomonas* reads to MAG completeness and ability to retrieve functional data. Furthermore, we defined the bacterial communities associated with bacterial spot that could provide insight into future studies to define community members related to disease. Lastly, we determined if MAGs created by our pipeline and whole genome isolates from the same sample were comparable.

A culture independent approach is a strength of MGS; however, the relationship between pathogen read abundance and isolation is not well-established. While all but three of our samples with *Xanthomonas* read abundance greater than 0.1% had successful isolation, there were cases where read abundance and successful isolation were not consistent. Discrepancies between metagenomic and culture-dependent methods in plant pathology have been reported before with *Pectobacterium* in potato (Seaman et al. 2025). However, we showed our metagenomic approach, regardless of *Xanthomonas* read abundance, was possible for species identification via ANI. While our *in silico* experiment demonstrated lower read abundance corresponded with lower BUSCO scores and MAG coverage, ANI values were not affected by read abundance. For example, sample 12T3 had 25,660 *Xanthomonas* reads, for a *Xanthomonas* read abundance of 0.149%, yet had an ANI of 96.80% with the *Xep* reference genome (Supplemental Table 5).

Taxonomic identification is essential for novel pathogen identification, microbial surveillance, or pathogens that pose a biosecurity threat (Ko et al. 2022; Thomas et al. 2017; Venbrux et al. 2023), and our work emphasizes the importance of ANI in MGS diagnostics. Future work is still needed to determine the lower limits of identification for plant pathogens.

A subset of samples (122T, 29T2, 2Roma, and 25T2bindweed) had the highest ANI value with *X. arboricola*, traditionally known as the causal agent of bacterial canker on fruit trees (Zarei et al. 2022). *X. arboricola* has been previously associated with bacterial spot disease symptoms (Mbega et al. 2012; Myung et al. 2010) but can be known a commensal pathogen (Pena et al. 2024). The presence of *X. arboricola* and the association with disease on tomato in Ohio is not currently described. We are now aiming to elucidate the role of *X. arboricola* in Ohio’s fresh market tomato and pepper production systems.

Precise microbial function and virulence factor identification in metagenomic data associated with disease depends on genome completeness, which inherently relies on abundance of pathogenic reads present. We found that the percentage of *Xanthomonas* reads in a given sample corresponds to MAG BUSCO score completeness, supported by both our *in silico* experiment and field samples. Since T3Es mediate both virulence and host recognition (Grant et al. 2006; White et al. 2009), these virulence factors represent important markers for surveillance. In contrast to species designations via ANI, T3E presence, determined by blastx, required a higher threshold for identification. A set of 36 effectors was identified across all *Xee* MAGs (where percent *Xanthomonas* was equal to or greater than 3%) and whole genomes from both *in silico* and field samples. Although it has been reported that at least 45 variable T3SS effectors make up the bacterial spot effector repertoire, the species and pathovar T3SS effector vary for each bacterial spot species (Potnis et al. 2011; Schwartz et al. 2015). For example, (Schwartz et al. 2015) identified only 23 effectors belonging to *Xhg*, while identified 29 and 30 effectors belonging to *Xep* and *Xee*, respectively. Therefore, a threshold based on abundance of *Xanthomonas*, rather than a set number of effectors, is the best approach. Notably, MAGs derived from samples where *Xanthomonas* comprised 3% or more of reads recovered the same expected 36 effectors from WGS.

Some effectors are of particular interest for disease management. A specific example is the recent emergence of race 4 *Xep*, which carries a nonfunctional *avrXv3* effector that is recognized by the tomato *Xv3* resistance gene (Bernal et al. 2022). Race 4 *Xep* can therefore overcome disease resistance and is a concern in production systems, making the tracking and presence/absence of T3SS effectors remains relevant for management decisions. We determined that in all WGS/MGS pairs, *avrXv3* was not present, however, no pairs clustered with *Xep* reference genomes. Additional whole genome sequencing of all *Xanthomonas* field isolates recovered can provide additional T3SS effector identification.

Prior studies have revealed a core set of bacterial spot and species/strain specific effectors (Potnis et al. 2011; Roach et al. 2019; Schwartz et al. 2015). Potnis et al. (2011) determined a core set of 11 T3Es among the four bacterial spot species and pathovars, and Schwartz et al. (2015) added two more effectors to this list. From our MAG/WGS comparison, one or both genomes contained the core set, with one exception: *xopR*. *xopR* was absent from all of our pairings, which is notable as *xopR* is among the nine core effectors of all *Xanthomonas* spp. (Ryan et al. 2011) and is known to disrupt plant defense response (Akimoto-Tomiyama et al. 2012). We identified effectors present in all pairings (identified by at least one in our pairs) not previously known to be present in *Xee* and *Xhg*, which were *xopAM*, *xopAQ* and *xopAZ* (Potnis et al. 2011; Schwartz et al. 2015). Interestingly, *xopAZ* has only recently been identified from bacterial spot pathogens in Taiwan (Chen et al. 2024), and previously, was found in *X. arboricola* (Assis et al. 2021). Further studies need to be conducted using phylogenetics to define if these effectors are present from a minority community member, not resolved by the MAG or potentially horizontally transferred. We expect more work is needed to determine if these thresholds are true for tracking functions in the plant microbiome.

There were minor effector identification discrepancies between MAGs and whole genome pairs from our field samples. While we hypothesized whole genomes may identify more effectors than their MAG counterparts, we did not anticipate MAGs identifying unique effectors compared to their whole genomes. Despite this, we suspect this could be explained by our assignment-first approach. MAG creation was based on Kraken2’s taxonomic classification from our bacterial spot genome database. Since we used the –include_children tag in conjunction with the *Xanthomonas* taxid (#338) using the extraction_kraken tool, we believe some reads could be mis-recovered. However, we support our use of this approach, as it still allowed us to identify

### Xanthomonas species

Researchers have provided evidence that the environment plays a strong role in affecting the microbiome over the host in humans (Rothschild et al. 2018) and plants (Bokulich et al. 2014). In our study, bacterial community composition is significantly impacted by host but not region (Supplemental Figure 1A, 1B) and moreover Shannon diversity is not impacted by region or host. Therefore, more work is needed to investigate the drivers of bacterial spot community composition. Our work also highlighted other taxa that could potentially play a role in symptom development. Abundance of varying *Pseudomonas* species in our samples was observed including *P*. *viridflava*, *P. syringae*, and *P. cichorii*, which have been associated with causing disease on tomato (Jones et al. 1984; Schneider and Grogan 1977; Timilsina et al. 2017). We found a high abundance of both *P. viridflava* and *P. syringae* in our samples, highlighting the potential relationship between bacterial spot xanthomonads and other tomato pathogens but did not observe any abundance of *P. cichorii*. The presence of *P. fulva* was not as surprising as we initially thought. *P. fulva* was found to be associated with bacterial spot in another study, and notably, is related to human infection and plant pathogen inhibition (Adeniji et al. 2020; Liu et al. 2014; Newberry et al. 2020). This microbiome analysis reveals that there is compounding evidence *Pseudomonas* is associated with bacterial spot, but further investigation is needed to determine the overall impact of their presence and role in symptom development. The abundance of *Xanthomonas* and *Stenotrophomonas maltophilia* in our bacterial spot samples was expected, as it is known to be associated with both plants and bacterial spot (Berg and Martinez 2015; Newberry et al. 2020). Notably, *S. maltophilia* has also been associated with necrotic spots on tomato seeds in Bulgaria (Stoyanova et al. 2018), but is primarily known for being an emergent, multi-drug resistant pathogen in humans, especially in hospitals (Brooke 2012).

Overall, we provide an approach for bacterial spot diagnostics with MGS and an assignment first approach, where pathogen lineage and function were determined. Our results suggest growers viewed Ohio PathID as a potentially valuable and compatible diagnostic approach, without prior interaction with the diagnostic results. These favorable perceptions may reflect the fact growers were not required to pay, implement, or assume risk with the technology, instead receiving its benefits. We demonstrated MAGs coming from samples with a high abundance of pathogen and their whole genome counterparts are comparable and provide consistent information. However, the microbial community findings rival traditional plant pathology techniques such as Koch’s postulates, where a singular causal agent is isolated, identified, symptoms are reproduced, and isolated once again. Despite its importance in pathogen identification, this traditional technique limits our understanding of plant health to a singular pathogen, singular host framework. As we observed other potential pathogens present in the taxa described, there could be additional microorganism contributing to symptomology. An inquiry for the future is how much do these additional community members contribute to the disease beyond the agent who can alone produce the symptoms. Metagenomics in plant disease research creates pathways to hypothesize and test synergistic relationships between microbial community members.

## Acknowledgements

We would like to thank the extension agents and growers who participated in this project.

---

**Supplemental Figure 1.**
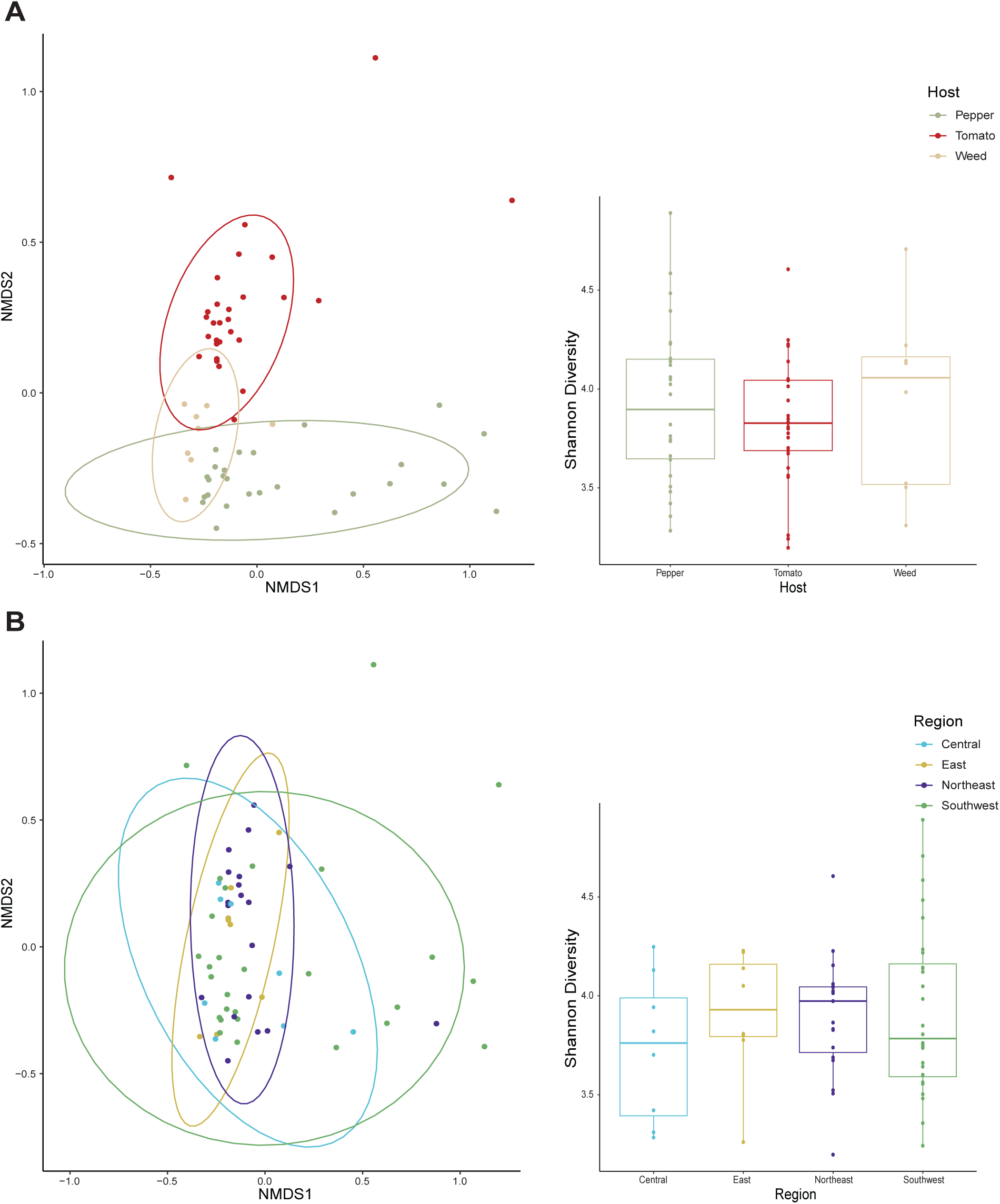
Bray-Curtis ordination plot of overall community composition and Shannon diversity of samples by host and region. **A)** PERMANOVA analysis indicates host significantly affects community composition (R^2^ = 0.33, P = 0.001), however, there is no significant difference in Shannon diversity across samples (Kruskal-Wallis, P = 0.79). **B)** Region does not significantly impact community composition (R^2^ = 0.06, P = 0.23) nor Shannon diversity (Kruskal-Wallis, P = 0.72).

## Notes

### Competing Interest Statement

The authors have declared no competing interest.

