## Supplemental Table 1 for "Metagenomics for bacterial spot pathogen and virulence factor tracking for Ohio fresh market tomato and pepper production"

ANI

| | Xe_85-10 | Xhg_SM775-1 | Xp_SM1806-1 | Xv_LM159 | rep1-005 | rep1-01 | rep1-02 | rep1-03 | rep1-04 | rep1-05 | rep1-06 | rep1-07 | rep1-08 | rep1-09 | rep1-1 | rep1-125 | rep1-150 | rep1-175 | rep1-2 | rep1-250 | rep1-3 | rep1-4 |
|---|---|---|---|---|---|---|---|---|---|---|---|---|---|---|---|---|---|---|---|---|---|---|
| Xe_85-10 | 1 | 0.88106336 | 0.98704783 | 0.87994987 | 0.9980542 | 0.99748221 | 0.99758604 | 0.99758882 | 0.99734489 | 0.99744693 | 0.99738665 | 0.99741174 | 0.99750517 | 0.99768006 | 0.99769393 | 0.99785645 | 0.99805867 | 0.998240641 | 0.99827848 | 0.99851728 | 0.99862105 | 0.9986779 |
| Xhg_SM775-1 | 0.8810298 | 1 | 0.88112926 | 0.88145342 | 0.91749865 | 0.91045174 | 0.91075893 | 0.9071776 | 0.9051628 | 0.90221406 | 0.89994153 | 0.89800692 | 0.8958516 | 0.89397138 | 0.8926187 | 0.88916676 | 0.88670159 | 0.884961468 | 0.88341902 | 0.88222867 | 0.88162403 | 0.8812057 |
| Xp_SM1806-1 | 0.98706142 | 0.88114751 | 1 | 0.88079287 | 0.98914793 | 0.98799496 | 0.98733057 | 0.98697454 | 0.98679846 | 0.98670914 | 0.98661972 | 0.98658783 | 0.98653847 | 0.98645175 | 0.98634026 | 0.98633864 | 0.98636921 | 0.986537344 | 0.98641331 | 0.98652354 | 0.98648496 | 0.98644466 |
| Xv_LM159 | 0.87991941 | 0.8815512 | 0.8807724 | 1 | 0.91627383 | 0.91094726 | 0.90812099 | 0.90553838 | 0.90364128 | 0.90088616 | 0.89920909 | 0.89703047 | 0.89514967 | 0.89327258 | 0.89149598 | 0.88799591 | 0.8856626 | 0.883779251 | 0.8821584 | 0.88092182 | 0.88014345 | 0.87976195 |
| rep1-005 | 0.99805976 | 0.91771641 | 0.98924767 | 0.91627383 | 1 | 0.99994187 | 0.9995539 | 0.99953623 | 0.99949832 | 0.99918115 | 0.99897895 | 0.99900604 | 0.99893782 | 0.99890021 | 0.99892366 | 0.99890452 | 0.9984718 | 0.998907951 | 0.99887669 | 0.99894465 | 0.99887387 | 0.99889019 |
| rep1-01 | 0.99748489 | 0.91045174 | 0.98803665 | 0.91094726 | 0.99994187 | 1 | 0.99954665 | 0.99940563 | 0.99897264 | 0.99893395 | 0.99885604 | 0.99886243 | 0.99883093 | 0.99881047 | 0.99867921 | 0.99862714 | 0.99835243 | 0.998541206 | 0.99854846 | 0.99856079 | 0.99847006 | 0.99852861 |
| rep1-02 | 0.99758776 | 0.91091712 | 0.9873449 | 0.90816726 | 0.9995539 | 0.99954665 | 1 | 0.99954874 | 0.99925078 | 0.99907256 | 0.99900829 | 0.99893065 | 0.99881219 | 0.9987709 | 0.99872004 | 0.99859893 | 0.99845049 | 0.998600389 | 0.99862239 | 0.99858231 | 0.99846503 | 0.99852086 |
| rep1-03 | 0.9975874 | 0.90725137 | 0.98693185 | 0.90556881 | 0.99953623 | 0.99940563 | 0.99954874 | 1 | 0.99956671 | 0.99929549 | 0.9991524 | 0.99892135 | 0.99878692 | 0.99867874 | 0.9986158 | 0.99851661 | 0.99843595 | 0.998507189 | 0.99847549 | 0.99847303 | 0.99843804 | 0.99842933 |
| rep1-04 | 0.99738411 | 0.90523787 | 0.98684643 | 0.90371209 | 0.99949832 | 0.99910763 | 0.99925078 | 0.99957323 | 1 | 0.99963066 | 0.99941862 | 0.9991837 | 0.99899669 | 0.99892268 | 0.99879641 | 0.9986394 | 0.99847993 | 0.998482484 | 0.99846595 | 0.99842757 | 0.99837373 | 0.99835055 |
| rep1-05 | 0.99747823 | 0.90231487 | 0.98672581 | 0.90094877 | 0.99913848 | 0.99889324 | 0.99908249 | 0.99932158 | 0.99964029 | 1 | 0.999705 | 0.99939553 | 0.99920151 | 0.99903327 | 0.99891253 | 0.9987228 | 0.99858605 | 0.998591699 | 0.99854769 | 0.9985398 | 0.9984862 | 0.99843399 |
| rep1-06 | 0.99742107 | 0.89999965 | 0.98669577 | 0.89928983 | 0.99897895 | 0.99885604 | 0.99901198 | 0.99916104 | 0.99941102 | 0.99969906 | 1 | 0.99964899 | 0.99936872 | 0.99919933 | 0.99901032 | 0.99878829 | 0.99862618 | 0.998549008 | 0.99853369 | 0.9984933 | 0.99842856 | 0.99843084 |
| rep1-07 | 0.99747252 | 0.89808756 | 0.98661629 | 0.89702921 | 0.99900604 | 0.99886243 | 0.99893433 | 0.99892992 | 0.99918322 | 0.99940545 | 0.99965596 | 1 | 0.99967505 | 0.99947109 | 0.99924705 | 0.99896687 | 0.99875137 | 0.998686392 | 0.99866795 | 0.99859931 | 0.99853627 | 0.99853056 |
| rep1-08 | 0.99754502 | 0.89596258 | 0.98655699 | 0.89516607 | 0.99893782 | 0.99883769 | 0.99881838 | 0.99878731 | 0.99898962 | 0.99920185 | 0.9993757 | 0.99966515 | 1 | 0.99969823 | 0.99941543 | 0.99906727 | 0.99886087 | 0.998763501 | 0.99872945 | 0.99869241 | 0.99864702 | 0.99864369 |
| rep1-09 | 0.99767009 | 0.89407995 | 0.98645611 | 0.89329277 | 0.99890021 | 0.99881721 | 0.99877342 | 0.99867864 | 0.99892128 | 0.9990329 | 0.99922227 | 0.99946059 | 0.99969771 | 1 | 0.99971024 | 0.99931258 | 0.99900855 | 0.998886198 | 0.99883052 | 0.99875933 | 0.9987428 | 0.9987369 |
| rep1-1 | 0.99767328 | 0.8927542 | 0.9863229 | 0.89153465 | 0.99892366 | 0.99868591 | 0.99872256 | 0.99863813 | 0.99879509 | 0.99891207 | 0.99900931 | 0.99923018 | 0.99940913 | 0.99970483 | 1 | 0.99945088 | 0.99913596 | 0.998990307 | 0.99889025 | 0.99877801 | 0.99874466 | 0.99873266 |
| rep1-125 | 0.99787132 | 0.88925526 | 0.9863914 | 0.88803652 | 0.99890452 | 0.99862935 | 0.99863417 | 0.99853133 | 0.99862643 | 0.99873015 | 0.99879281 | 0.99897179 | 0.99908436 | 0.99931723 | 0.99946001 | 1 | 0.99950477 | 0.999295201 | 0.9991527 | 0.99900969 | 0.9989312 | 0.99895162 |
| rep1-150 | 0.99806181 | 0.88684065 | 0.98641797 | 0.88566115 | 0.9984718 | 0.99833201 | 0.99843263 | 0.99844488 | 0.99847311 | 0.99859755 | 0.99862167 | 0.99875393 | 0.9988758 | 0.99901125 | 0.99912455 | 0.99950581 | 1 | 0.999623501 | 0.99939777 | 0.99925468 | 0.99919952 | 0.99914678 |
| rep1-175 | 0.99826685 | 0.88506649 | 0.98649974 | 0.88378199 | 0.99890795 | 0.99851423 | 0.99853048 | 0.99851685 | 0.99846285 | 0.99858918 | 0.99854854 | 0.99868209 | 0.99880299 | 0.99891142 | 0.99899077 | 0.99928766 | 0.9996232 | 1 | 0.99969346 | 0.99948735 | 0.99940113 | 0.99939033 |
| rep1-2 | 0.99829488 | 0.88352966 | 0.98636482 | 0.88219211 | 0.99887669 | 0.99851821 | 0.99857856 | 0.99846103 | 0.99845436 | 0.99855976 | 0.99853163 | 0.99865948 | 0.99873956 | 0.9988464 | 0.99889911 | 0.99914472 | 0.99939824 | 0.999693272 | 1 | 0.99964383 | 0.9994972 | 0.99944247 |
| rep1-250 | 0.99851573 | 0.8823149 | 0.98650015 | 0.88099576 | 0.99894465 | 0.99853982 | 0.99854197 | 0.99845906 | 0.9984382 | 0.99853971 | 0.99851317 | 0.99859775 | 0.99869385 | 0.99876827 | 0.99877801 | 0.99899122 | 0.99924086 | 0.999487724 | 0.99964242 | 1 | 0.99978772 | 0.99971201 |
| rep1-3 | 0.99862055 | 0.88175239 | 0.98645908 | 0.88017857 | 0.99887387 | 0.99847927 | 0.99846847 | 0.99843876 | 0.99834548 | 0.99846767 | 0.99843018 | 0.99851944 | 0.99863464 | 0.99875282 | 0.99875443 | 0.99895639 | 0.99918583 | 0.999402042 | 0.9994844 | 0.99978774 | 1 | 0.999791 |
| rep1-4 | 0.99866609 | 0.88128558 | 0.98642724 | 0.87979137 | 0.99889019 | 0.99855713 | 0.99852445 | 0.99842777 | 0.99835044 | 0.99843801 | 0.99843164 | 0.99853207 | 0.9986436 | 0.99875952 | 0.99875584 | 0.99895314 | 0.99915711 | 0.999393103 | 0.99944247 | 0.99971123 | 0.99980392 | 1 |

Coverage

| | Xe_85-10 | Xhg_SM775- | Xp_SM1806- | Xv_LM159 | rep1-005 | rep1-01 | rep1-02 | rep1-03 | rep1-04 | rep1-05 | rep1-06 | rep1-07 | rep1-08 | rep1-09 | rep1-1 | rep1-125 | rep1-150 | rep1-175 | rep1-2 | rep1-250 | rep1-3 | rep1-4 |
|---|---|---|---|---|---|---|---|---|---|---|---|---|---|---|---|---|---|---|---|---|---|---|
| Xe_85-10 | 1 | 0.62498249 | 0.85423058 | 0.63282114 | 0.02121115 | 0.05349222 | 0.14278416 | 0.24793832 | 0.35455125 | 0.45329341 | 0.5420361 | 0.61743434 | 0.67966033 | 0.7287746 | 0.77060556 | 0.83735789 | 0.87353743 | 0.894023808 | 0.90488574 | 0.91314739 | 0.91606345 | 0.91773722 |
| Xhg_SM775- | 0.64582498 | 1 | 0.65368313 | 0.65904315 | 0.00779657 | 0.02001724 | 0.0590504 | 0.10847815 | 0.16168026 | 0.2135907 | 0.26782558 | 0.31591534 | 0.36483538 | 0.40532551 | 0.44239056 | 0.5146902 | 0.560886 | 0.593899293 | 0.61559408 | 0.6355587 | 0.64414304 | 0.64958161 |
| Xp_SM1806- | 0.89864339 | 0.66448614 | 1 | 0.6692233 | 0.02067442 | 0.05175866 | 0.13859098 | 0.24115144 | 0.34556467 | 0.44318137 | 0.52955079 | 0.60342655 | 0.66476167 | 0.71363652 | 0.75413625 | 0.82070871 | 0.85638352 | 0.876895995 | 0.88785066 | 0.89678558 | 0.89999559 | 0.90138527 |
| Xv_LM159 | 0.63530074 | 0.64067884 | 0.63893873 | 1 | 0.00723724 | 0.01912972 | 0.05623564 | 0.10208087 | 0.15312121 | 0.20427593 | 0.2546305 | 0.30093105 | 0.34495959 | 0.3840357 | 0.4221815 | 0.49452345 | 0.54086164 | 0.575366664 | 0.59748717 | 0.61884125 | 0.62927394 | 0.63530964 |
| rep1-005 | 0.94987336 | 0.33652557 | 0.88015462 | 0.32303375 | 1 | 0.99671396 | 0.98339596 | 0.98042445 | 0.97283427 | 0.96977172 | 0.9721721 | 0.97327297 | 0.97303293 | 0.97759366 | 0.98353668 | 0.98758422 | 0.98924794 | 0.990522622 | 0.99236843 | 0.99358518 | 0.99049779 | 0.99049779 |
| rep1-01 | 0.96505004 | 0.34956958 | 0.88687486 | 0.34300668 | 0.40116 | 1 | 0.98429234 | 0.97968498 | 0.97497102 | 0.97493104 | 0.97589382 | 0.97823248 | 0.98007476 | 0.98334955 | 0.98742388 | 0.98834668 | 0.99162147 | 0.994246632 | 0.99470304 | 0.99594566 | 0.99510947 | 0.99476967 |
| rep1-02 | 0.96326585 | 0.38488334 | 0.89159167 | 0.37822994 | 0.14832498 | 0.36897461 | 1 | 0.98977831 | 0.98157299 | 0.97934687 | 0.97972018 | 0.98187639 | 0.98320981 | 0.98511257 | 0.98678809 | 0.99096316 | 0.99470999 | 0.996480403 | 0.99709967 | 0.99788 | 0.99760158 | 0.99784005 |
| rep1-03 | 0.96799797 | 0.41025221 | 0.89640579 | 0.39642273 | 0.08549592 | 0.21220487 | 0.57163219 | 1 | 0.98735218 | 0.98371925 | 0.98290024 | 0.98501698 | 0.98389373 | 0.98476969 | 0.98747762 | 0.99163613 | 0.99380406 | 0.995685047 | 0.99703541 | 0.99802457 | 0.99818895 | 0.99771672 |
| rep1-04 | 0.9721085 | 0.42873256 | 0.90165755 | 0.41727848 | 0.05945239 | 0.14829276 | 0.39774144 | 0.69252108 | 1 | 0.9916415 | 0.98826129 | 0.98798323 | 0.98652417 | 0.98623651 | 0.98874158 | 0.9918963 | 0.9939494 | 0.995951437 | 0.99709958 | 0.99796561 | 0.99829776 | 0.99796763 |
| rep1-05 | 0.97243687 | 0.4429916 | 0.90469909 | 0.43586345 | 0.04633752 | 0.11598175 | 0.31028382 | 0.5393665 | 0.77500416 | 1 | 0.99341813 | 0.99070491 | 0.98835099 | 0.98764739 | 0.98924354 | 0.9917868 | 0.9938988 | 0.99583964 | 0.99677211 | 0.99744172 | 0.99774095 | 0.99739824 |
| rep1-06 | 0.97396698 | 0.46453524 | 0.90482349 | 0.45369418 | 0.03883293 | 0.09692519 | 0.25959795 | 0.45062473 | 0.64582199 | 0.83046864 | 1 | 0.99444069 | 0.99063039 | 0.98880603 | 0.98981817 | 0.99181039 | 0.99355147 | 0.995962689 | 0.99677194 | 0.9975412 | 0.99825528 | 0.99772459 |
| rep1-07 | 0.97353373 | 0.48089457 | 0.90522802 | 0.47145012 | 0.03414024 | 0.08530229 | 0.22838133 | 0.39658079 | 0.56664587 | 0.72670554 | 0.87229354 | 1 | 0.99442168 | 0.99089462 | 0.99145675 | 0.99201568 | 0.99375398 | 0.995881232 | 0.99654343 | 0.99767609 | 0.99835742 | 0.99783533 |
| rep1-08 | 0.97274956 | 0.50391474 | 0.90472986 | 0.49025676 | 0.03095748 | 0.07753208 | 0.20749095 | 0.35931609 | 0.51325218 | 0.65738938 | 0.78786756 | 0.90139639 | 1 | 0.99412257 | 0.99308288 | 0.99189272 | 0.99381266 | 0.995814687 | 0.99671626 | 0.9977128 | 0.99817029 | 0.99775699 |
| rep1-09 | 0.97381195 | 0.52244583 | 0.90636439 | 0.5089921 | 0.02901094 | 0.07253324 | 0.19387156 | 0.33557493 | 0.47860101 | 0.61269696 | 0.73351665 | 0.83761889 | 0.92676011 | 1 | 0.99645844 | 0.99334984 | 0.99440404 | 0.996449353 | 0.99729099 | 0.99814221 | 0.99837121 | 0.99787467 |
| rep1-1 | 0.97393918 | 0.53979063 | 0.90632576 | 0.5295599 | 0.02763063 | 0.0690315 | 0.18381059 | 0.31840169 | 0.45414181 | 0.58076605 | 0.69480949 | 0.79269057 | 0.87582458 | 0.94236938 | 1 | 0.99419872 | 0.99476776 | 0.996409138 | 0.99719149 | 0.99797361 | 0.99817227 | 0.99753817 |
| rep1-125 | 0.97348042 | 0.57751917 | 0.90735048 | 0.57044957 | 0.02556372 | 0.06358156 | 0.16991727 | 0.29413587 | 0.41916644 | 0.53565354 | 0.64028064 | 0.7294054 | 0.80407539 | 0.86303268 | 0.91322843 | 1 | 0.996611 | 0.996738848 | 0.99708177 | 0.99796109 | 0.99789118 | 0.99751875 |
| rep1-150 | 0.97383535 | 0.60413219 | 0.90773189 | 0.59902638 | 0.02456008 | 0.06118328 | 0.16358711 | 0.28282842 | 0.40309384 | 0.51508225 | 0.61532791 | 0.70083744 | 0.77253897 | 0.8284017 | 0.87581841 | 0.95470556 | 1 | 0.99828487 | 0.99746464 | 0.99808961 | 0.99802295 | 0.99750853 |
| rep1-175 | 0.97381099 | 0.62524341 | 0.9077248 | 0.62280996 | 0.02404785 | 0.0599815 | 0.16020683 | 0.27694788 | 0.39459492 | 0.50427418 | 0.60261838 | 0.68607356 | 0.75628166 | 0.81093735 | 0.85690875 | 0.93240017 | 0.97438578 | 1 | 0.99876363 | 0.99860224 | 0.9980461 | 0.99770688 |
| rep1-2 | 0.9734557 | 0.64043624 | 0.90849637 | 0.63980089 | 0.02382253 | 0.05934571 | 0.15848536 | 0.27423694 | 0.39071387 | 0.49933829 | 0.59652949 | 0.67891074 | 0.74837834 | 0.80237704 | 0.84788949 | 0.92183463 | 0.96203754 | 0.986780678 | 1 | 0.99889464 | 0.99831897 | 0.99751274 |
| rep1-250 | 0.97344978 | 0.65487799 | 0.90823967 | 0.65569811 | 0.02364305 | 0.05888511 | 0.15718087 | 0.27218441 | 0.38779508 | 0.49537036 | 0.59182448 | 0.67392333 | 0.74255657 | 0.79583345 | 0.84090459 | 0.91433479 | 0.95374328 | 0.977100443 | 0.98904487 | 1 | 0.99847293 | 0.99775773 |
| rep1-3 | 0.97358962 | 0.66184982 | 0.90890156 | 0.6649743 | 0.0234982 | 0.05869727 | 0.1567976 | 0.27162613 | 0.38690357 | 0.49423248 | 0.59073147 | 0.67269876 | 0.74112525 | 0.79402187 | 0.83902662 | 0.91188785 | 0.9510449 | 0.97380584 | 0.98579782 | 0.99534357 | 1 | 0.99834122 |
| rep1-4 | 0.97236921 | 0.66578135 | 0.90772074 | 0.66960527 | 0.02342798 | 0.05845304 | 0.1563803 | 0.27071078 | 0.38560217 | 0.49266879 | 0.58877841 | 0.67055113 | 0.73883054 | 0.79141369 | 0.83628291 | 0.90905645 | 0.94806887 | 0.970645226 | 0.98213087 | 0.99182116 | 0.99543785 | 1 |

| BUSCO scores | |
|---|---|
| Xe_85-10 | 99.1% |
| Xhg_SM775-12 | 99.3% |
| Xp_SM1806-17 | 99.2% |
| Xv_LM159 | 99.6% |
| rep1-005 | 0.1% |
| rep1-01 | 0.3% |
| rep1-02 | 0.8% |
| rep1-03 | 2.0% |
| rep1-04 | 4.2% |
| rep1-05 | 6.3% |
| rep1-06 | 9.6% |
| rep1-07 | 14.5% |
| rep1-08 | 19.0% |
| rep1-09 | 26.1% |
| rep1-1 | 31.6% |
| rep1-125 | 44.9% |
| rep1-150 | 58.0% |
| rep1-175 | 69.8% |
| rep1-2 | 80.4% |
| rep1-250 | 91.1% |
| rep1-3 | 95.5% |
| rep1-4 | 98.6% |
