## Supplemental Table 2 for "Metagenomics for bacterial spot pathogen and virulence factor tracking for Ohio fresh market tomato and pepper production"

| Farm Number | County | Varieties | Sequencing Name | Xantho reads | Xantho reads/total reads (%) | Xanthomonas isolated? |
|---|---|---|---|---|---|---|
| 1 | Geauga | Tomato: Roma, Mariana | 1MT | 298192 | 2.230935724 | yes |
| 2 | Geauga | Pepper: Bell, California Wonder | 2CW | 2710 | 0.020049477 | no |
| 2 | Geauga | Tomato: Roma | 2Roma | 13878 | 0.109843907 | no |
| 3 | Geauga | Pepper: Bell, Red Knight | 3RKP | 544 | 0.004295867 | no |
| 3 | Geauga | Tomato: Mountain Fresh | 3MFT | 107334 | 0.794192584 | no |
| 4 | Geauga | Pepper: Bell, PS 0994-1819 | 41819 | 2100 | 0.014120013 | no |
| 5 | Geauga | Pepper: Bell, Galileo | 5Galileo | 2110 | 0.013440177 | no |
| 6 | Franklin | Tomato: Mountain Fresh or New Girl | 6MFT | 1252 | 0.009532436 | no |
| 7 | Franklin | Tomato: Martha Washington | 7MWT | 312 | 0.002176269 | no |
| 7 | Franklin | Pepper: Bell, Ace | 7BAP | 104 | 7.29998E-05 | no |
| 8 | Morrow | Pepper: Bell, PS 0994-1819 | 81819 | 369126 | 3.071551201 | no |
| 8 | Morrow | Pigweed: By Pepper | 8Pig | 32516 | 0.233140913 | no |
| 9 | Crawford | Pepper: Bell, PS 0994-1819 | 91819 | 5228 | 0.030124252 | no |
| 10 | Huron | Pepper: Serrano, Altiplano | 10SAP | 4946 | 0.032163792 | no |
| 11 | Erie | Pepper: Bell, PS 0994-1819 | 111819 | 490238 | 3.547263587 | yes |
| 12 | Erie | Tomato 2: Cherry, Indigo Blue Berries | 122T | 83406 | 0.645703608 | yes |
| 12 | Erie | Tomato 3: Cherry, Sweet 100 | 12T3 | 25660 | 0.148617808 | yes |
| 13 | Cuyahoga | Tomato: Wapsipinicon Peach | 13WPT | 1568 | 0.00961772 | no |
| 13 | Cuyahoga | Tomato: Currant, White Currant | 13WCT | 9246 | 0.066411374 | no |
| 14 | Butler | Pepper: Jalapeno, Early Girl | 14EGJP | 154 | 0.001056199 | no |
| 14 | Butler | Pigweed: By Jalapeno, Early Girl Peppers | 14EGJPPig | 792 | 0.005484395 | no |
| 15 | Pickaway | Tomato: Field, 3 rows in (variety unknown) | 15T3 | 6478 | 0.045018437 | no |
| 15 | Pickaway | Pigweed: By Field, 3 rows in tomatoes | 15T3Pig | 836 | 0.005652738 | no |
| 16 | Mahoning | Tomato 1: Roma, Plum Royal | 16PRT | 369602 | 2.539839043 | yes |
| 16 | Mahoning | Tomato 2: Unknown variety | 16T2 | 380478 | 2.183810446 | yes |
| 17 | Columbiana | Tomato 1: Roma, Mariana | 17MRT | 46152 | 0.356436855 | yes |
| 18 | Belmont | Tomato 2: Unknown variety | 18T2 | 1234 | 0.007324286 | no |
| 19 | Belmont | Tomato: Primo Red | 19PRT | 20676 | 0.115515311 | no |
| 20 | Monroe | Pepper: Sweet Banana, Sweet Savannah | 20SSP | 114 | 0.000845239 | no |
| 21 | Champaign | Tomato: New Girl 2 (other side of field) | 21NG2T | 3668 | 0.026438063 | no |
| 21 | Champaign | Pepper: Bell, Sprinter | 21SBP | 244 | 0.00147483 | no |
| 22 | Champaign | Tomato: Brandywine | 22Brandy | 10916 | 0.066618365 | yes |
| 23 | Greene | Pepper: Ghost | 23GhostP | 2908 | 0.016620387 | no |
| 23 | Greene | Pepper: Carmen | 23CarP | 720 | 0.005509815 | no |
| 23 | Greene | Pepper Fruit with Lesion: PS 0994-1819 | 231819Fruit | 900 | 0.006289286 | no |
| 24 | Tuscarawas | Pepper: Jalapeno | 24JP | 14294 | 0.096312837 | yes |
| 24 | Tuscarawas | Tomato: Mountain Fresh | 24MFT | 812 | 0.006390399 | no |
| 25 | Belmont | Tomato 2: Unknown variety | 25T2 | 620 | 0.005667877 | no |
| 25 | Belmont | Bindweed: By Tomato 2 tomatoes | 25T2Bindweed | 14244 | 0.105525072 | yes |
| 25 | Belmont | Tomato Fruit with Lesions: By Tomatoes in Block 3 (no leaves from this area) | 25T3Fruit | 354164 | 1.53517215 | yes |
| 26 | Clinton | Tomato: Garden Peach | 26GPT | 74194 | 0.383495896 | no |
| 26 | Clinton | Pigweed: By Garden Peach tomatoes | 26GPTPig | 1770 | 0.012143128 | no |
| 26 | Clinton | Pepper: Escamillo | 26EP | 716 | 0.004521485 | no |
| 27 | Scioto | Tomato 3: Cherry | 27T3 | 222050 | 1.321969338 | yes |
| 27 | Scioto | Weed: By Tomato 3 Cherry tomatoes (spikey weed) | 27T3Weed | 956 | 0.005626189 | no |
| 27 | Scioto | Pepper 6: Orange spaceships, unknown variety | 27P6 | 539688 | 3.938233236 | yes |
| 27 | Scioto | Pepper 6 Fruit with Lesion: Orange spaceship | 27P6Fruit | 246738 | 1.999863184 | yes |
| 27 | Scioto | Pepper 7: Carolina Reaper | 27P7 | 147338 | 0.950795197 | yes |
| 27 | Scioto | Pepper 8: Ghost | 27P8 | 268752 | 2.438720735 | yes |
| 27 | Scioto | Healthy Tomato 1 | 27HT1 | 18260 | 0.143845062 | no |
| 27 | Scioto | Healthy Pepper 1 | 27HP1 | 714 | 0.005067791 | no |
| 28 | Clark | Tomato 1: Beefsteak (unknown variety) | 28T1 | 232246 | 1.665162199 | yes |
| 28 | Clark | Healthy Tomato 1 | 28HT1 | 242 | 0.001705198 | no |
| 28 | Clark | Pepper 1: Bell (unknown variety) | 28P1 | 306990 | 2.585064443 | yes |
| 28 | Clark | Pepper 2: Hot, red, small (unknown variety) | 28P2 | 205868 | 1.612875173 | yes |
| 28 | Clark | Healthy Pepper 2 | 28HP2 | 256 | 0.001815001 | no |
| 28 | Clark | Pepper Fruit with Lesions: Pepper 4 | 28P4Fruit | 240978 | 2.010088395 | yes |
| 29 | Medina | Tomato 2: Beefsteak (unknown variety) | 29T2 | 45640 | 0.379570081 | yes |
| 30 | Medina | Tomato 2: Roma (unknown variety) | 30T2 | 4340 | 0.029041488 | no |
| 31 | Ashland | Pigweed: By Tomato 1 tomatoes | 31T1Pig | 514 | 0.004526906 | no |
| 32 | Highland | Tomato 2: Orange, pointed, globular tomato (unknown variety) | 32T2 | 20600 | 0.156050652 | yes |
| 32 | Highland | Pigweed: By Tomato 2 tomatoes | 32T2Pig | 4872 | 0.05308744 | no |
| 32 | Highland | Tomato Fruit with Lesions: Tomato 2 | 32T2Fruit | 77556 | 0.54850301 | yes |
