## Supplemental Table 4 for "Metagenomics for bacterial spot pathogen and virulence factor tracking for Ohio fresh market tomato and pepper production"

| Assembly Accession | Organism | Strain |
| --- | --- | --- |
| GCA_003993395.1 | Xanthomonas arboricola | BRIP62410 |
| GCA_002939805.1 | Xanthomonas arboricola | CFBP6827 |
| GCA_001854165.1 | Xanthomonas euvesicatoria | 85-10 |
| GCA_020879845.1 | Xanthomonas euvesicatoria | DC 96-5 |
| GCA_001908795.1 | Xanthomonas euvesicatoria | LMG930 |
| GCA_001009625.1 | Xanthomonas gardneri | SM775-12 |
| GCA_001908775.1 | Xanthomonas gardneri | ICMP7383 |
| GCA_001009485.1 | Xanthomonas perforans | GEV872 |
| GCA_001908855.1 | Xanthomonas perforans | LH3 |
| GCA_021608005.1 | Xanthomonas perforans | SM1806-17 |
| GCA_001908815.1 | Xanthomonas vesicatoria | LM159 |
| GCA_001908725.1 | Xanthomonas vesicatoria | LMG911 |
| GCA_017724035.1 | Xanthomonas euvesicatoria pv. alfalfae | |
| GCA_040529065.1 | Xanthomonas sacchari | |
| GCA_900183975.1 | Xanthomonas fragariae | |
| GCA_009769165.1 | Xanthomonas hyacinthi | |
| GCA_001908725.1 | Xanthomonas vesicatoria | |
| GCA_014236795.1 | Xanthomonas theicola | |
| GCA_000961215.1 | Xanthomonas citri pv. citri | |
| GCA_008370835.2 | Xanthomonas oryzae pv. oryzicola | |
| GCA_013388375.1 | Xanthomonas campestris pv. raphani | |
| GCA_905367715.1 | Xanthomonas arboricola pv. juglandis | |
| GCA_017301775.1 | Xanthomonas translucens pv. undulosa | |
| GCA_008639345.1 | Xanthomonas phaseoli pv. dieffenbachiae | |
| GCA_041519315.1 | Xanthomonas axonopodis pv. maculifoliigardeniae | |
| GCA_000277895.2 | Xanthomonas vasicola pv. musacearum | |
| GCA_025266575.1 | Xanthomonas prunicola | |
| GCA_009883735.1 | Xanthomonas cucurbitae | |
| GCA_021906995.1 | Xanthomonas cerealis pv. cerealis | |
| GCA_040616535.1 | Xanthomonas cannabis | |
| GCA_030168915.1 | Xanthomonas graminis pv. graminis | |
| GCA_032698475.1 | Xanthomonas dyei | |
| GCA_032697525.1 | Xanthomonas hydrangeae | |
| GCA_000454545.1 | Xanthomonas cassavae | |
| GCA_035027565.3 | Xanthomonas campestris pv. campestris | |
| GCA_040202155.2 | Xanthomonas hortorum pv. hederae | |
| GCA_000019585.2 | Xanthomonas oryzae pv. oryzae | |
